# FastSK: Fast Sequence Analysis with Gapped String Kernels

**DOI:** 10.1101/2020.04.21.053975

**Authors:** Derrick Blakely, Eamon Collins, Ritambhara Singh, Andrew Norton, Jack Lanchantin, Yanjun Qi

## Abstract

Gapped *k*-mer kernels with Support Vector Machines (gkm-SVMs) have achieved strong predictive performance on regulatory DNA sequences on modestly-sized training sets. However, existing gkm-SVM algorithms suffer from slow kernel computation time, as they depend exponentially on the sub-sequence feature-length, number of mismatch positions, and the task’s alphabet size. In this work, we introduce a fast and scalable algorithm for calculating gapped *k*-mer string kernels. Our method, named FastSK, uses a simplified kernel formulation that decomposes the kernel calculation into a set of independent counting operations over the possible mismatch positions. This simplified decomposition allows us to devise a fast Monte Carlo approximation that rapidly converges. FastSK can scale to much greater feature lengths, allows us to consider more mismatches, and is performant on a variety of sequence analysis tasks. On 10 DNA transcription factor binding site (TFBS) prediction datasets, FastSK consistently matches or outperforms the state-of-the-art gkmSVM-2.0 algorithms in AUC, while achieving average speedups in kernel computation of ∼100× and speedups of ∼800× for large feature lengths. We further show that FastSK outperforms character-level recurrent and convolutional neural networks across all 10 TFBS tasks. We then extend FastSK to 7 English-language medical named entity recognition datasets and 10 protein remote homology detection datasets. FastSK consistently matches or outperforms these baselines. Our algorithm is available as a Python package and as C++ source code^1^.

## 1 Introduction

String kernels in conjunction with Support Vector Machines (SK-SVM) achieve strong prediction performance across a variety of sequence analysis tasks, with widespread use in bioinformatics and natural language processing (NLP). SK-SVMs are a popular technique for DNA regulatory element identification [22, 10, 25, 7, 19, 27, 8], and bio-medical named entity recognition [23, 15]. SK-SVMs are also popular baselines for evaluating the quality of deep learning models [3, 10, 25] for analyzing variant impacts.

The key to the success of string kernel methods is their use of simple, yet expressive, substring features to compute a similarity function between sequences. In turn, the similarity function defines an inner product space, where an SVM classifier can be trained. The approach easily enables comparison of arbitrary length sequences, obviates sequence alignment issues, captures task-relevant pattern information, and is simpler than other pattern detection tools, such as position-weight matrices [28, 2]. Viewed as a type of “feature engineering”, string kernels yield simpler and lower variance models than deep learning. One consequence is that they show strong performance without consuming vast amounts of training data (for example, see Figure 4).

**Figure 1:**
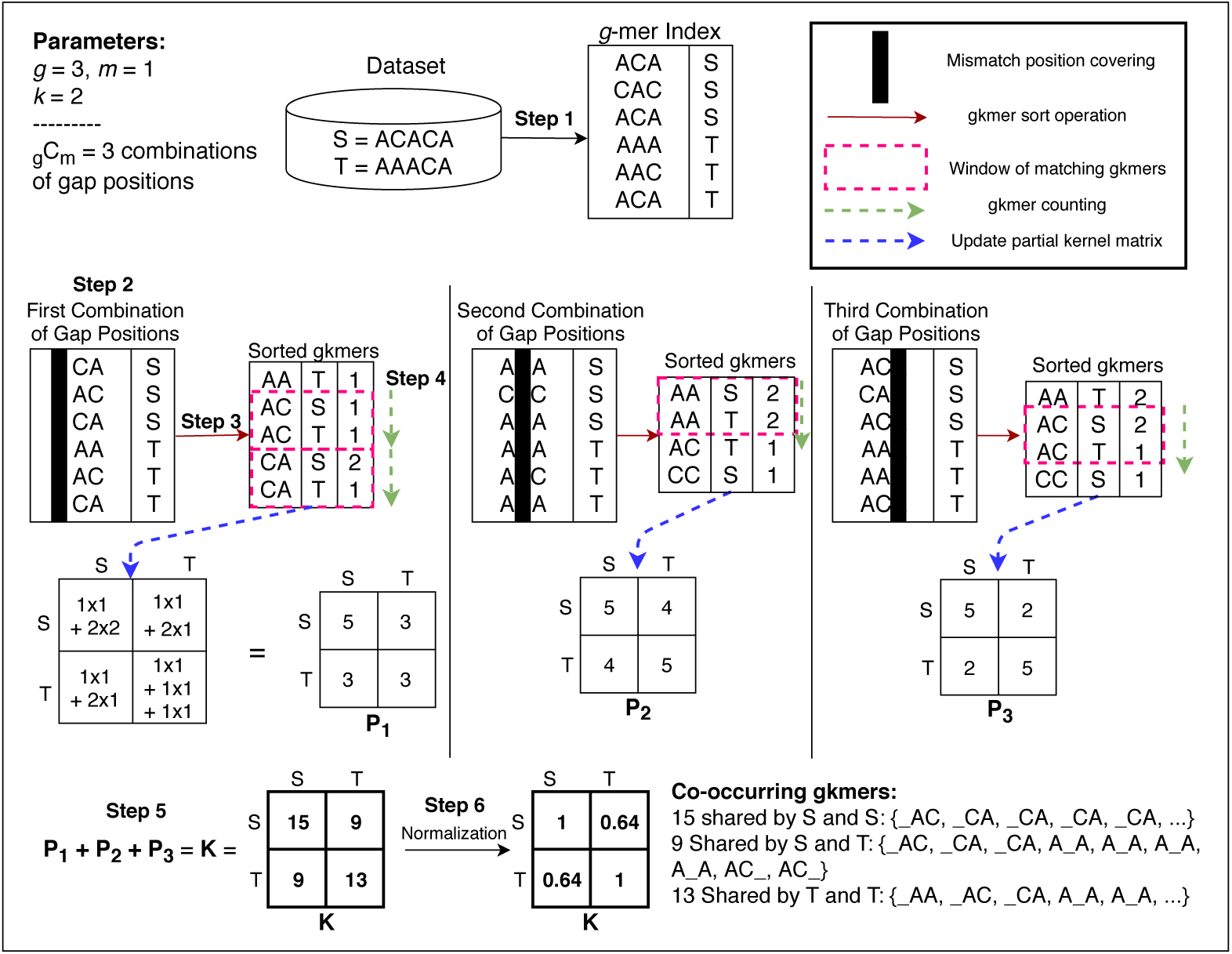
A demonstration of the FastSK -Exact algorithm for *g* = 3, *m* = 1, and *k* = 2 using a small dataset with only two sequences (S and T). (1) All *g*-mers are extracted from the dataset and stored in a table, along with the IDs of the sequences from which they were extracted. (2) One of the possible 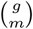 combinations of mismatch positions is removed to produce the gkmers. (3) The gkmers are sorted lexicographically. (4) Co-occurrences of the gkmers are counted and stored in the corresponding *partial kernel matrix P*_*i*_. Steps (2-4) are repeated until each *P*_*i*_ is computed. (5) Once all partial matrices are computed, they are summed to produce the unnormalized kernel matrix *K*. (6) Kernel values are normalized.

**Figure 2:**
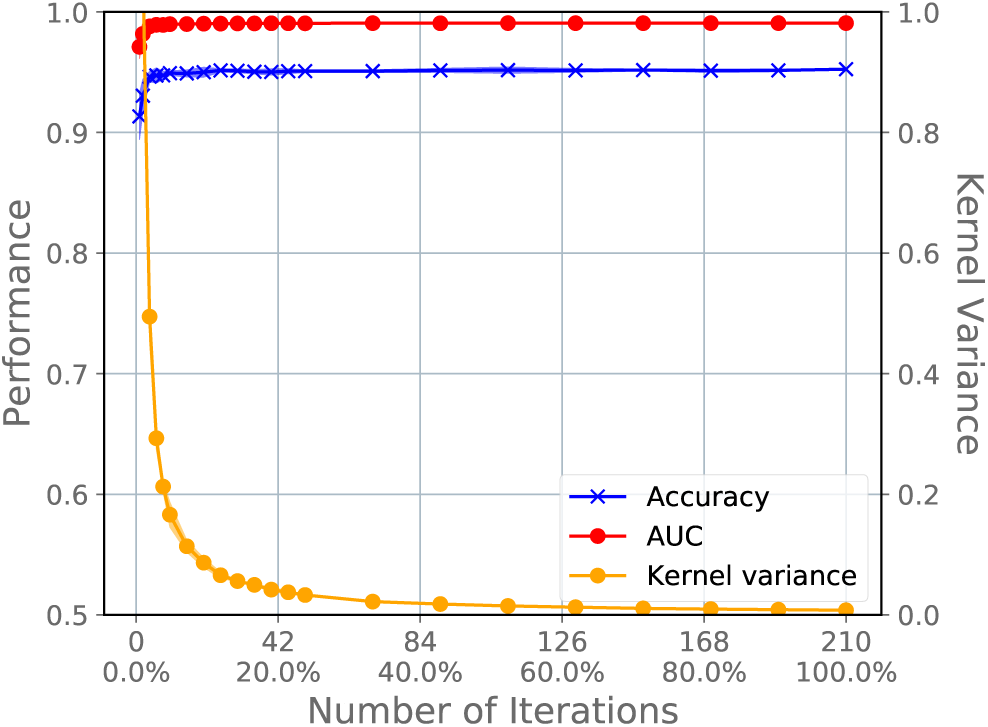
Only a small proportion of the possible mismatch combinations are needed for strong performance. We use the EP300 TFBS (DNA) dataset with *g* = 10 and *m* = 4, hence 210 possible mismatch combinations. On the x-axis, we show the number of mismatch positions used and the percentage of the total possible below each one. The left y-axis shows the performance metric (AUC and accuracy) on the test set, while the right y-axis shows the variance of the kernel matrix. Together, these show that the FastSK approximation algorithm converges rapidly, which corresponds to strong performance.

**Figure 3:**
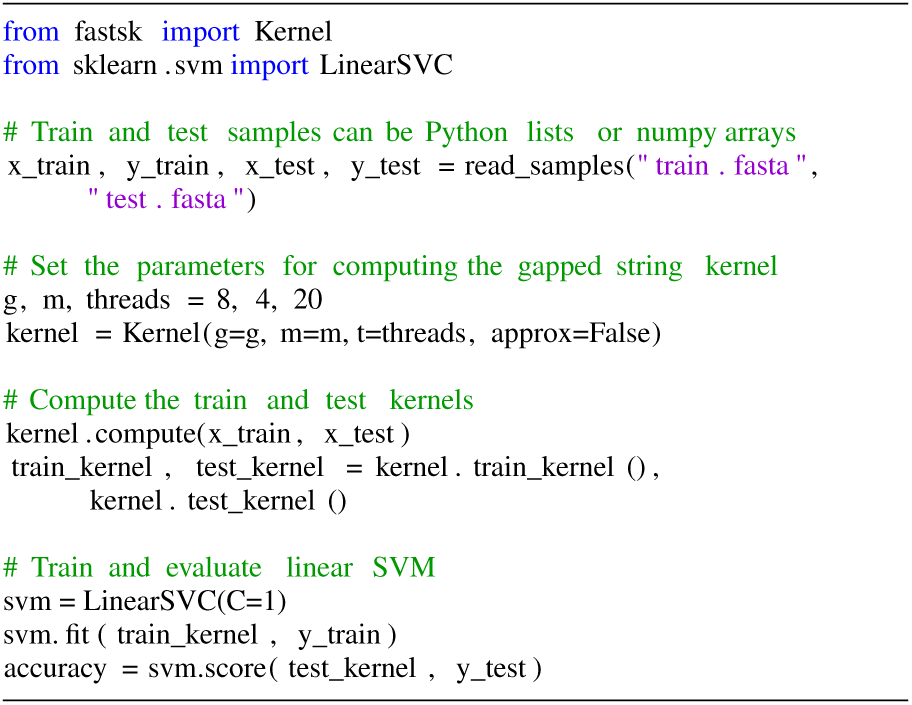
FastSK is easy to use in Python in conjunction with the Scikit-Learn library. Here, we show an example of computing a gapped string kernel and then using the train and test kernels to train and evaluate a linear SVM. Training many other models (kernel SVM, logistic regression, etc.) is straightforward as well.

**Figure 4:**
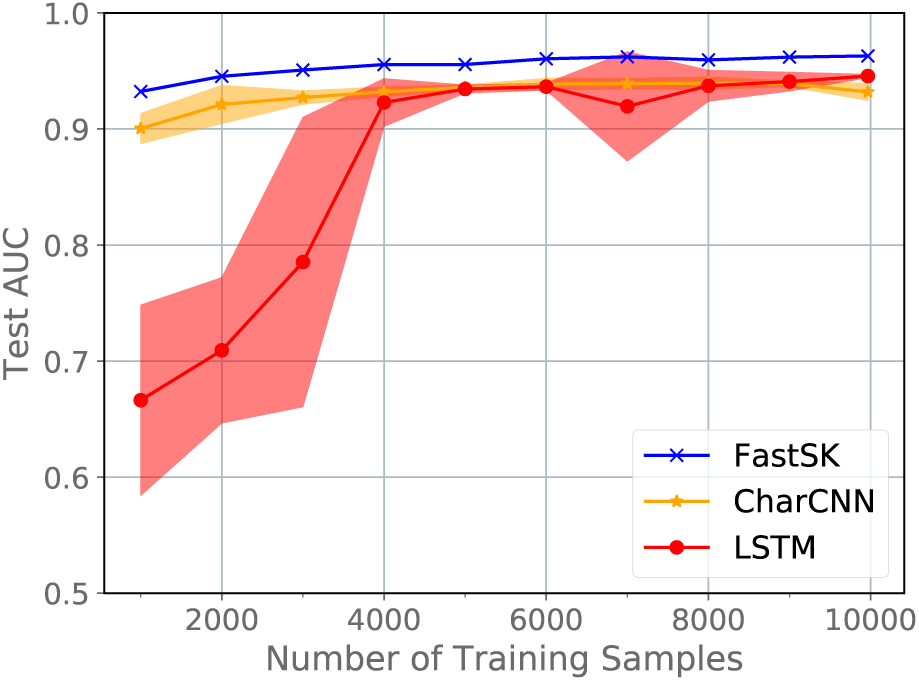
We vary the proportion of training samples used for a DNA sequence classification task (TFBS prediction on the ZZZ3 ENCODE ChIP-Seq dataset). Training sizes vary from 1,000 to 9,996 samples (the maximum). Each point is the average of 5 runs, with the shaded region showing standard error.

In greater detail, string kernels use substring features to map sequences to fixed-dimension feature vectors. Such a mapping is referred to as a *feature map*. For example, one of the most popular string kernels is the *k*-spectrum kernel [21], which maps sequences to vectors using contiguous length-*k* substrings, or *k*-mers. In this vector space, the *i*-th component of a sequence’s feature vector is simply the number of times the *i*-th possible *k*-mer occurs in that sequence. The *kernel function* then takes a pair of sequences and returns their similarity score—the inner product of their feature vectors in the vector space induced by the feature map. Importantly, a kernel function serves a “kernel trick” in the Support Vector Machine framework. In turn, this provides several benefits of its own: SVMs learn an optimal separating hyperplane to classify sequences and show excellent generalization [29]. Moreover, they obtain excellent prediction performance without needing as many training samples as state-of-the-art deep learning models (shown by Figure 4).

The spectrum string kernel and derivatives such as the Gapped K-Mer kernels via Support Vector Machine (gkm-SVMs) [7] and (*k, m*)-mismatch kernel [4] are responsible for many of the successes of string kernels. However, a major problem for spectrum kernels is that the dimensionality of the feature space is |∑|^*k*^, where |∑| is the alphabet of characters appearing in the sequences. This presents three major problems. First, kernel computation becomes infeasible for even modest values of |∑| or *k*. For example, spectrum kernels are infeasible for roughly *k >* 10, yet many transcription factor binding sites are up to 20 basepairs long. Furthermore, spectrum methods scale poorly to large alphabets, such as protein or natural language [4, 20, 30]. The second challenge is that as *k* increases, the odds of observing any particular *k*-mer within a sequence rapidly goes to zero. Therefore, the feature vectors are both extremely large and extremely sparse, which makes models trained on these feature vectors highly prone to overfitting and poor generalization. Third, existing algorithms that overcome these challenges leave much to be desired; for example, popular trie-based approaches (e.g., [7, 4]) still exhibit exponential dependence on |∑| [20, 27], *k*, and *m*. On the other hand, counting-based methods rely on complex “mismatch statistics” to indirectly obtain feature counts [17, 6, 27], however still fail to scale to greater feature lengths.

Together, these issues present major limitations to the practical utility of *k*-mer string kernel methods. To solve these problems, we introduce a novel string kernel algorithm called FastSK. It makes four high-level contributions:

1. We take inspiration from [7] and use *gapped k-mer* features, or *gkmers* for short, which are a more compact feature set that greatly reduces the size of the feature space and risk of overfitting. These gapped *k*-mer features consist of an overall length of *g* = *k* + *m*, length-*k* non-contiguous substrings within the features, and *m* mismatch positions.
2. We decompose the kernel function into a set of 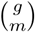 independent counting operations to count the gapped *k*-mer features shared between sequences. This formulation is simpler and faster than the state-of-the-art string kernel algorithms. Furthermore, the independence of the counting operations mean that the algorithm is naturally parallelizable. We exploit this advantage to create a fast multithreaded implementation.
3. We take advantage of the independence of each mismatch position to create a scalable Monte Carlo approximation algorithm. This approach randomly samples the possible mismatch positions until the variance of the kernel matrix converges. We show the approximation algorithm, called FastSK -Approx, converges rapidly irrespective of the parameters *g* and *m*. Therefore, FastSK -Approx allows scaling to greater feature lengths.
4. Empirically, we show that FastSK matches or outperforms state-of-the-art string kernel methods, convolutional neural networks (CNN) and LSTM models in test performance across 10 DNA TFBS datasets, 10 protein remote homology detection datasets, and 7 medical named entity recognition tasks.

FastSK provides a list of benefits compared to previous works: (1) It scales to greater feature lengths, effectively running in 𝒪(1) time with respect to the feature length. (2) It does not show AUC decay at greater feature lengths. (3) Its running time is independent of the alphabet size |∑|, allowing it to generalize to any sequence analysis task. (4) It outperforms two deep learning models when using modestly-sized training sets. (5) Finally, FastSK has a nice bonus property: it is a much simpler algorithm because it *directly* computes the kernel function instead of using complicated “mismatch statistics.” (6) Most importantly, we make our algorithm available as a toolkit via C++ source code in Github and via a Python package.

## 2 Background

### 2.1 Support Vector Machines

Support Vector Machines learn a linear predictive model *f*(*x*) = *ŷ* = **x · w** + *b*. In the case of linearly-separable data, SVMs optimize the parameters **w** by learning a pair of max-margin hyperplanes given by:

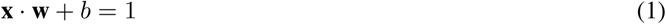

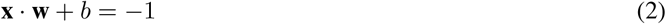

This is achieved by minimizing ∥**w**∥_2_; because we want to maximize the distance between the planes, which is 2*/*∥**w**∥_2_, we minimize ∥**w**∥_2_. To keep training points from being inside the margin, we also impose the constraint:

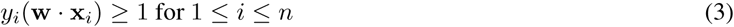

where *n* is the number of training samples.

A non-linear, or kernelized, SVM follows roughly this same structure, but uses a kernel function *K*(·, ·) to compute the pair-wise similarities between samples. As such, a string kernel function *K* is easily applied in the Support Vector Machine (SVM) framework [29]. In this case, the predicted class of a sample *x* is given by

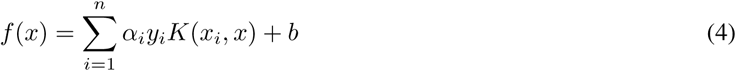

where *x*_*i*_ and *y*_*i*_ are the *i*th training sample and its label, respectively. Each *α*_*i*_ is a weight, where if *α*_*i*_ ≠ 0, *α*_*i*_ corresponds to a *support vector* and *b* is a learned additive bias.

### 2.2 String Kernels

String kernel methods compare arbitrary-length sequences by mapping them to a fixed inner product space. The key component is the feature map *ϕ* : 𝒮 →ℝ^*p*^, where is the set of all strings composed from the alphabet ∑. The dimensionality *p* of the vector space depends on the particular string kernel’s feature map. The canonical example is the spectrum kernel by [21], which uses simple length-*k* substrings, or *k*-mers. Given a string *x* = (*s*_1_, *s*_2_, …, *s*_|*x*|_) with each *s*_*i*_ ∈ ∑ and a substring size *k*, the spectrum kernel *ϕ*_*S*_ maps *x* to a vector indexed by all possible length-*k* substrings from ∑^*k*^. Under this feature map, the *i*th dimension of the vector *ϕ*_*S*_ (*x*) holds the number of times the *i*th possible *k*-mer ∈ ∑^*k*^ occurs in *x*. The spectrum kernel function *K*_*S*_ then provides a similarity score of two sequences *x* and *y* as the inner product of their spectrum feature vectors:

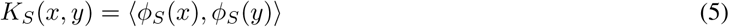

A string kernel function *K* is easily applied in the Support Vector Machine (SVM) framework [29]. Importantly, the inner product of a string kernel can be defined *implicitly* as part of the kernel function. That is, without invoking the explicit mapping of strings to their full feature vectors in the vector space. For example, the spectrum kernel is also given by

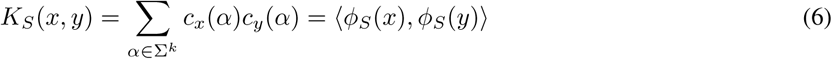

where *c*_*x*_(*α*) and *c*_*y*_(*α*) return the counts of *k*-mer *α* in sequences *x* and *y*, respectively. This view provides an important intuition: the feature function *ϕ*(·) can be evaluated *implicitly*; that is, without fully mapping the sequences to their feature vectors. Another important intuition is that a *k*-mer *α* only contributes to *K*_*S*_ (*x, y*) if it is present in *both* sequences; that is, both *c*_*x*_(*α*) and *c*_*y*_(*α*) have to be non-zero. Therefore, evaluation of *K*_*S*_ (*x, y*) reduces to counting the co-occurrences of each possible *α*. In fact, this view shows we must compare the *k*-mers between samples in order to determine if some *α* is present in both. Finally, to allow imprecise matching (e.g., permitting robustness to noise or single nucleotide polymorphisms), we can permit some number of *mismatches* when comparing the *k*-mers between sequences. This approach, called the (*k, m*)-mismatch kernel, defines the kernel function as

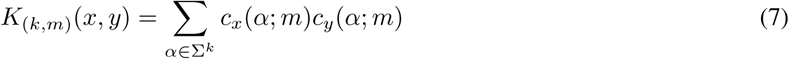

where *c*_*x*_(*α*; *m*) is the number of times the *k*-mer *α* appears in the sequence *x* with up to *m* mismatches. In the next section, we further elaborate upon this formulation.

### 2.3 (k,m)-mismatch Kernel

Since the introduction of the spectrum kernel, many string kernel variants and generalizations have been proposed, usually involving *mismatches* to incorporate noise into string comparisons. For example, the (*k, m*)-mismatch kernel from [20] retains the *k* parameter for substring lengths, while adding an *m* parameter to denote a number of *mismatches* permitted when comparing the *k*-mers of a pair of sequences.

A simple generalization of the *k*-spectrum kernel is the (*k, m*)-mismatch kernel [20], which permits up to *m mismatches* when determining if a pair of *k*-mers should contribute to the similarity of their respective sequences. Under the (*k, m*)-mismatch feature map, a string *x* is mapped to a |∑|^*k*^ -dimensional space by

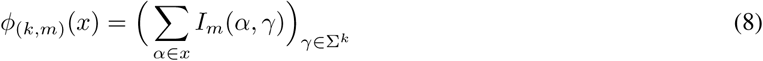

where *I*_*m*_(*α γ*) = 1 if the *k*mer *γ* is in the “mismatch neighborhood” of *α*, denoted by *N*_*k,m*_(*α*). The mismatch neighborhood *N*_*k,m*_(*α*) is simply the set of all *k*mers that differ from *α* by at most *m* characters. The *i*th index of *ϕ*_(*k,m*)_(*x*) is simply a count of how many times the *i*th possible *k*mer occurs in *x* if we allow up to *m* mismatches.

An influential approach by [17] uses this formulation to compute the kernel function *indirectly*. Intuitively, the similarity of sequences *x* and *y* is given by how many “neighboring” *k*-mers they share. Following this intuition, the trick is to compute the kernel function by counting how many *k*-mers from sequence *x* are contained in the mismatch neighborhoods of sequence *y*’s *k*-mers. As shown in [17], this reduces to counting the number of *k*-mers shared between *x* and *y* at each Hamming distance *d* ∈ ({0, 1, …, min(2*m, k*)} and then multiplying the counts (which the authors call “statistics”) by an appropriate combinatorial coefficient. We dub this approach and its derivatives “mismatch statistic string kernels.” The upside of the mismatch statistic approach is that it works well in the case of the (*k, m*)-mismatch kernel and is not computationally dependent on the alphabet size |∑ | (the feature space, however, is). The downside is that it has been applied to situations where it is not actually beneficial. We show this in section 3.5.

As for the (*k, m*)-mismatch kernel generally, there are significant shortcomings. First, because the feature space is of size |∑ | ^*k*^, operating in this space becomes deleterious for even moderately sized ∑ or *k*. Second, this is an extremely sparse feature space, as the probability of any particular *k*-mer appearing in a sequence quickly approaches 0 as *k* grows. As such, (*k, m*)-mismatch SK-SVMs are highly prone to overfitting. Third, most implementations use trie-based data structures, which also grow exponentially with ∑ and *k* [4, 20, 30]. Implementation that do not use tries often use a complex set of *mismatch statistics* to *indirectly* compute the co-occurrence counts [16, 17, 6].

Ultimately, none of these approaches are effective at meeting all of the following three criteria: (1) Feature set that is not exponential in the alphabet size |∑|. (2) Fast kernel computation algorithm that is scalable in ∑, and conceptually simple. (3) It should scale to greater feature lengths and numbers of mismatches. For example, efficiently handling transcription factor binding sites with 20 basepairs.

## 3 Proposed Algorithm: FastSK

### 3.1 Gapped k-mer Kernel

Like [7], we use a gapped *k*-mer, or gkmer, feature set. These features have three parameters: an overall feature length *g, m* mismatch (or gap^2^) positions, and *k* informative, non-mismatch positions (note that *k* + *m* = *g*). For example, as shown in figure 1, the string *S* = *ACACA*, contains the *g*-mer *g*_1_ = *ACA*, where *g* = 3. Now, the parameter *k* specifies a number of informative, non-mismatch positions within the *g*-mer. For example, if *k* = 2, then {*AC*_, *A*_*A*, _*CA*} are the set of possible *gapped k-mers* within *g*_1_. We use *m* (in this case *m* = 1) to denote the number of mismatch positions inside the *k*-mers. As with the *k*-spectrum and (*k, m*)-mismatch kernel, our kernel function is determined by the co-occurrences of *k*-mers, except in our case the *k*-mers are not contiguous features. The gapped *k*-mer string kernel function is given by

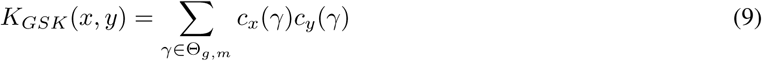

where Θ_*g,m*_ is the set of gapped *k*-mers with *m* mismatch positions appearing in the dataset; in contrast to the spectrum methods, we do not need to consider the entire feature space. Function *c*_*x*_(*γ*) gives the count of gapped *k*-mer *γ* in *x*.

### 3.2 Sort-and-Count Kernel Algorithm

We decompose equation 9 into a summation of multiple independent counting operations, where each operation handles a combination of mismatch positions. Our kernel function is given by:

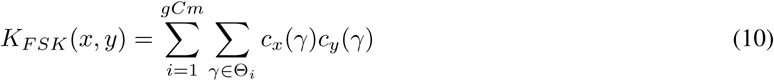

Here 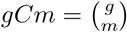, Θ_*i*_ denotes the set of gapped *k*-mers induced by the *i*th combination of *m* mismatch positions. Note that the sets Θ_*i*_ compose the full feature set Θ_*g,m*_. We give a demonstration of our algorithm for computing equation 10 in figure 1 and briefly summarize the algorithm here. First, we extractall *g*-mers from the dataset and store how many times each *g*-mer occurs in each sequence. Then for each of the 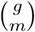 combinations of mismatch positions, we remove *m* mismatch positions from the *g*-mers to obtain a set of gkmers. Next, we sort the gkmers and count when they are shared in common between pairs of sequences. When we find that a gkmer *γ* occurs in both sequences *x* and *y*, we store the product *c*_*x*_(*γ*)*c*_*y*_(*γ*) in a *partial kernel matrix P*_*i*_. Denoting *P*_*i*_ as a function, the partial similarity score of *x* and *y* is given by:

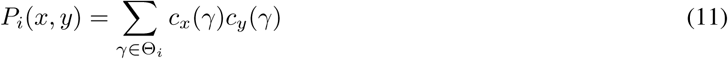

Importantly, we compute each *P*_*i*_ independently. This way the algorithm is both easy to parallelize and easy to approximate using random sampling. We demonstrate this experimentally in section 3.3. Ultimately, once each *P*_*i*_ for 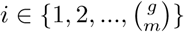 is computed, the full kernel matrix is given by

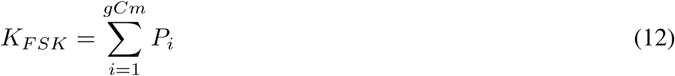

Where 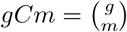. We show these steps in algorithm 0. Finally, we normalize the kernel matrix using

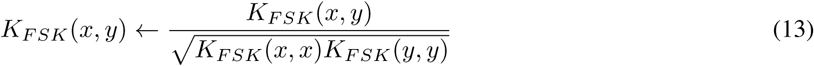

for pair of sequences (*x, y*). We refer to this algorithm as FastSK -Exact.

### 3.3 FastSK via fast Monte Carlo Approximation

Because FastSK -Exact runs with a coefficient of 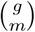, it is exponential in *g* and *m*. As such, it is unable to handle features roughly of size *g >* 15. However, many TF binding sites are up to 20 basepairs. If we let *k* = 6, we would have to compute more than 38,000 partial mismatch kernels. Moreover, even if *g* = 20 is not optimal, a thorough grid-search must include large values of *g* in the search space in order to rule them out. Therefore, there is a strong need to create gapped *k*-mer algorithms that can scale to greater feature lengths. To solve this problem, we introduce a Monte Carlo approximation algorithm called FastSK -Approx. FastSK -Approx is extremely fast even for large values of *g*, as it requires only a small random subset of the 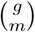 partial kernels *P*_*i*_. Empirically, we show that FastSK -Approx is roughly 𝒪(1) with respect to *g*.

To compute FastSK -Approx, we sample possible mismatch combinations for up to 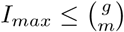 iterations. That is, at iteration 1 *≤ t ≤ I*_*max*_, we randomly sample (without replacement) a mismatch combination 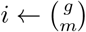 and compute the corresponding partial kernel matrix *P*_*i*_. We then compute the online mean kernel matrix 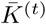 using *P*_*i*_ and 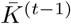. Furthermore, we compute a matrix of online standard deviations corresponding to the entries of 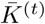 and use the average of these values, which we denote as *σ*^(*t*)^, to satisfy a convergence condition. Convergence is achieved when there is an approximately 95% probability that the online sample mean kernel 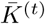 is within *δ* units of the true mean kernel matrix *µ*_*K*_. Here, *δ* is a user-determined parameter. In practice, we use *δ* = 0.025.

#### Algorithm: FastSK Exact

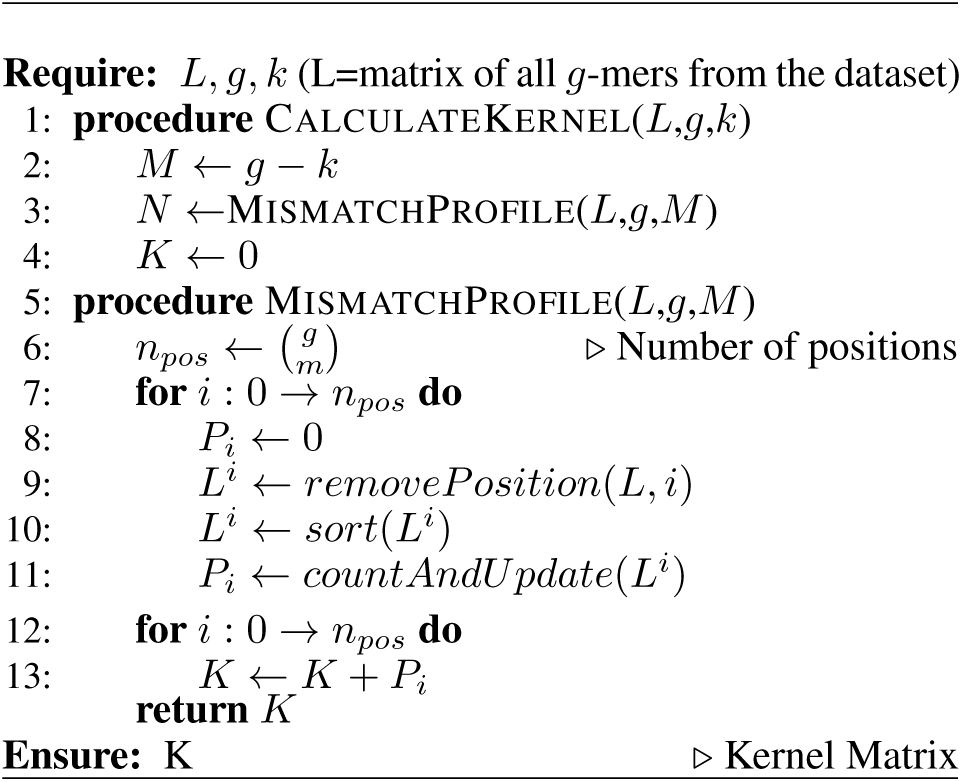

The idea rests on the Central Limit Theorem: we assume that for sufficiently large *t*, the sample mean kernel is normally distributed. Standardizing the variable 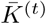, we have 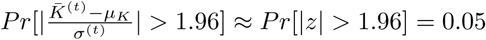, where 1.96 is the z-score for a 95% confidence interval. Therefore, the convergence condition is satisfied when

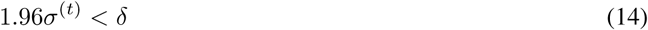

We show that FastSK -Approx converges rapidly, even for large values of *g* or *m*; it typically converges when *t* ≈ 50, which roughly corresponds to the number of samples needed to invoke the Central Limit Theorem. Furthermore, this means that FastSK -Approx is roughly 𝒪(1) with respect to *g*.

In Section 4.1.1 and Figure 6, we empirically show that FastSK -Approx converges rapidly across all datasets we tried. Our results show that FastSK -Approx consistently matches FastSK -Exact in prediction performance along with a significant speedup in time cost. Therefore FastSK -Approx is our default choice of FastSK.

**Figure 5:**
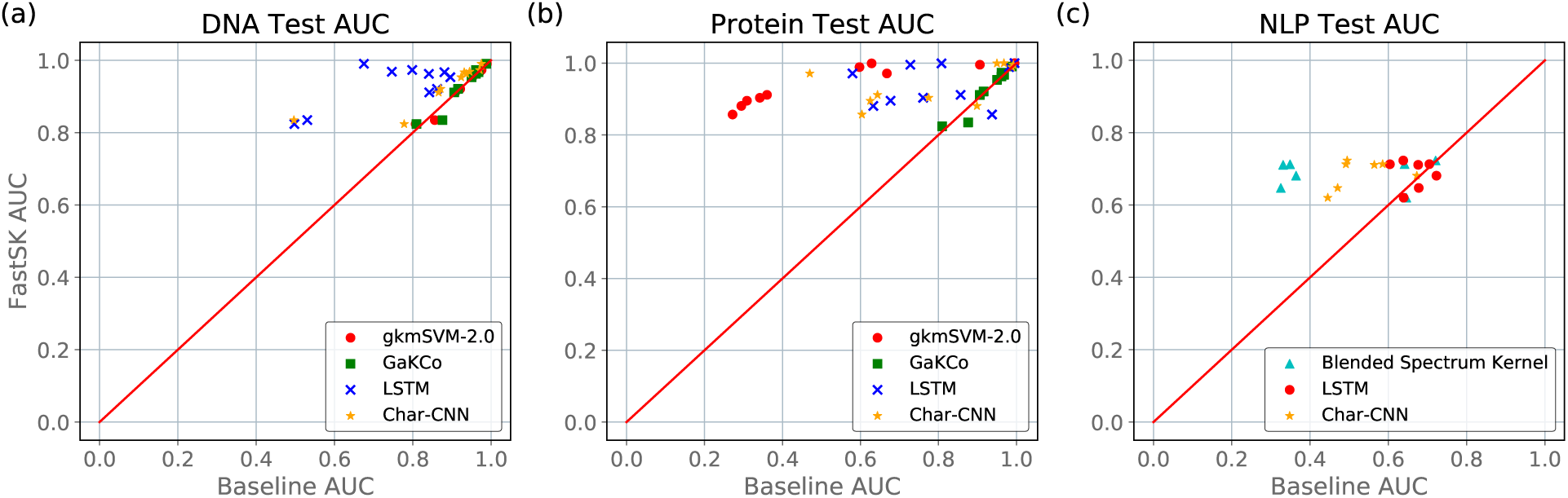
FastSK matches or outperforms the baselines across (a) DNA datasets, (b) protein datasets, and (c) medical NLP datasets. Please note that results in (b) relied on sequence information, therefore may be improved with more features like PSSM. In our context, experiments on 10 protein and 7 biomedical NLP datasets mainly serve to validate that FastSK can scale to sequence inputs with a larger alphabet.

**Figure 6:**
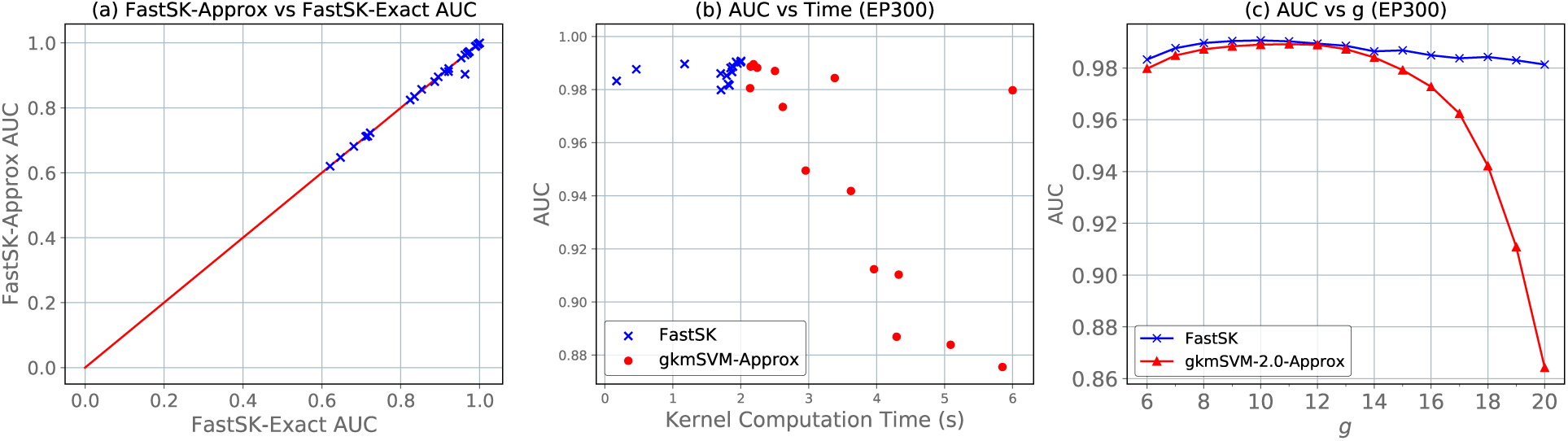
(a) FastSK -Approx with *I*_*max*_ = 50 achieves almost identical test AUC as FastSK -Exact. This is because very few partial kernels *P*_*i*_ are needed to achieve an excellent approximation of the kernel matrix. On average, the test AUC of FastSK -Approx is just 0.003 points lower than that of FastSK -Exact. (b) We plot AUC vs kernel computation time, where each point shows the AUC and kernel timing for a different value of *g* ∈ {6, …, 20}. I.e., 14 points for each FastSK and gkmSVM-2.0-Approx. We show that FastSK consistently retains both excellent AUC and low kernel computation times. On the other hand, gkmSVM-2.0-Approx degrades in AUC as the time needed to compute the kernel matrix increases. (c) The gkmSVM-2.0-Approx strategy results in severely degraded performance as *g* increases. It is therefore unreliable for searching the parameter space.

### 3.4 Software Details

Our algorithms are easily available as a PyPi package^3^ for Python. The package, called “fastsk”, consists of a well-optimized C++ implementation with a Python interface. We bind the C++ backend and Python interface using the Pybind11 library^4^ [13]. Therefore, we combine the simplicity and convenience of Python with the speed and low-level control of C++. To our knowledge, this is the only string kernel package that does so. We believe FastSK is the best string kernel toolbox for use within modern data science/machine learning toolboxes.

#### Usage

Unlike virtually all extant string kernel toolboxes, FastSK is designed to be easy to use in modern data science/machine learning workflows. For one, it is installed using a simple command (pip install fastsk). Two, kernel matrices can be computed in just a few lines of Python. Three, it is easy to use FastSK in conjunction with the standard classifiers in the Scikit-Learn library, including linear SVM (LinearSVC), kernel SVM (SVC), stochastic gradient descent (SGDClassifier), and logistic regression (LogisticRegression). Finally, evaluation and analysis are also quite simple, as metrics such as AUC and F1 score are trivial to compute using the Scikit-Learn metrics library. We illustrate a typical use-case in Figure 3.

#### Lower Triangular Kernel

Because a kernel matrix is symmetric by definition, we only store the lower triangle to save memory. In practice, the lower triangular kernel matrix is actually treated as a one-dimensional contiguous array in memory. This simplifies the code and improves cache utilization through better access locality.

### 3.5 Connecting to Related Work

#### Mismatch Statistic-Based String Kernels

While FastSK *directly* counts the gapped *k*-mers shared between sequences, previous works (e.g. [7, 8, 18, 19, 27]) *indirectly* compute the kernel function by inferring the counts from a set of *mismatch statistics*. These methods take inspiration from [17], which uses the notion of a *mismatch neighborhood* to efficiently compute the (*k, m*)-mismatch kernel. A mismatch neighborhood *N*_*d*_(*x, y*) is simply the number of pairs of *g*-mers from sequences *x* and *y* that have a Hamming distance ≤*d*. Using this “statistic,” the kernel function is inferred by multiplying *N*_*d*_(*x, y*) with an appropriate coefficient for each value of *d*. Given a value of *d*, the required coefficient is some kernel-dependent combinatorial value. Though the mismatch statistic idea was created to improve the efficiency of the (*k, m*)-mismatch kernel, we argue that it actually harms efficiency in the case of the gapped *k*-mer kernel.

To compute the gapped string kernel, gkmSVM [7] applies the mismatch neighborhood idea to gapped *k*-mers. The key observation is that a pair of *g*-mers with a Hamming distance of *d* share 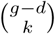 gkmers. Using this observation, the gkmSVM kernel is given by

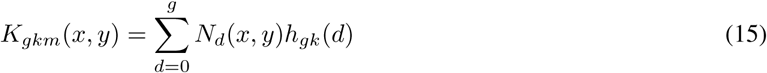

where

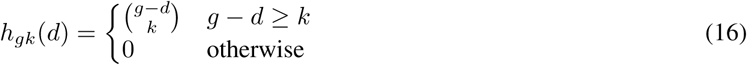

The key point is that this kernel function does not count the features of interest, which are gapped *k*-mers, or gkmers. Rather, it counts *g*-mers at various Hamming distances *d* and uses a coefficient *h*_*gk*_(*d*) to infer the gkmer counts.

##### Theorem 1. The FastSK kernel function is equivalent to the gkmSVM kernel function [7, 8].

*Proof*. Because *h*_*gk*_(*d*) = 0 for *d > m* = *g* − *k*, equation 15 is equal to

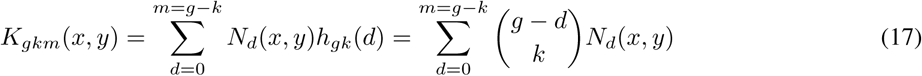

To *directly* count the gkmers, we replace 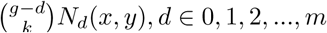 with 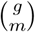, the number of possible mismatch positions. Therefore, at the *i*th mismatch position, we simply count the gapped *k*-mers in the set Θ_*i*_. Therefore, we have

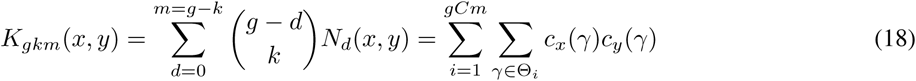

as desired. Our method is both faster and simpler because we cut out the middleman: we directly count the gkmers. It also allows us to create a better approximation algorithm than what is possible using equation 15, because we can sample mismatch positions to estimate the kernel function.

#### Trie-based Implementations

Several implementations, including gkmSVM and gkmSVM-2.0 [7, 8], use a *k*-mer tree (or trie) to compute the mismatch neighborhood *N*_*d*_(*x, y*) for each *d* 0, 1, 2, …, *g* and pair of sequences (*x, y*). These approaches have a branching factor equal to the alphabet size |∑ | (e.g., 4 for DNA) and depth of *g*. As such, they are impractical for tasks with larger alphabets, such as protein (∑ = 20) or English text (∑ ≥26) classification. Furthermore, they scale poorly with the parameter *g* and *m*.

#### Counting Implementations

GaKCo [27] is similar to FastSK in that it uses uses a sort-and-count algorithm. However, it differs from FastSK in that it follows the mismatch statistic formulation from [7]. The result is that GaKCo’s time complexity has a 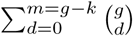 coefficient. In contrast, we simply have a coefficient of 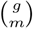.

## 4 Experimental Setup and Results

### Datasets

We evaluate FastSK using 10 transcription factor binding site (TFBS) DNA datasets derived from the ENCODE project, 10 protein remote homology datasets from the SCOP project, and 7 English-language medical named entity recognition datasets. Summary statistics and speedup results for our DNA, protein, and NLP datasets are shown in Tables 1, 2, and 3, respectively (tables 2 and 3 in appendix).

**Table 1:** 10 DNA Datasets (Data Statistics and Speedups)

| Dataset | Train | Test | Total | Avg Speedup× | Max Speedup× |
| --- | --- | --- | --- | --- | --- |
| CTCF | 2000 | 2000 | 4000 | 151 | 723 |
| EP300 | 2000 | 2000 | 4000 | 16 | 92 |
| JUND | 2000 | 2000 | 4000 | 151 | 757 |
| RAD21 | 2000 | 2000 | 4000 | 161 | 808 |
| SIN3A | 2000 | 2000 | 4000 | 16 | 88 |
| Pbde | 4500 | 5500 | 10000 | 18 | 107 |
| EP300_47848 | 6506 | 724 | 7230 | 162 | 833 |
| KAT2B | 6318 | 702 | 7020 | 161 | 809 |
| TP53 | 4432 | 494 | 4926 | 23 | 81 |
| ZZZ3 | 9966 | 1108 | 11074 | 18 | 111 |

**Table 2:**
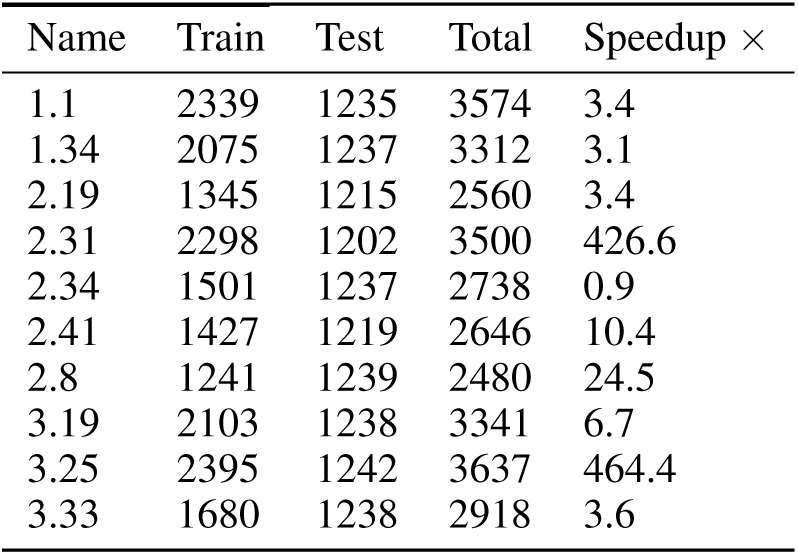
Protein Datasets

| Name | Train | Test | Total | Speedup × |
| --- | --- | --- | --- | --- |
| 1.1 | 2339 | 1235 | 3574 | 3.4 |
| 1.34 | 2075 | 1237 | 3312 | 3.1 |
| 2.19 | 1345 | 1215 | 2560 | 3.4 |
| 2.31 | 2298 | 1202 | 3500 | 426.6 |
| 2.34 | 1501 | 1237 | 2738 | 0.9 |
| 2.41 | 1427 | 1219 | 2646 | 10.4 |
| 2.8 | 1241 | 1239 | 2480 | 24.5 |
| 3.19 | 2103 | 1238 | 3341 | 6.7 |
| 3.25 | 2395 | 1242 | 3637 | 464.4 |
| 3.33 | 1680 | 1238 | 2918 | 3.6 |

**Table 3:** NLP Datasets

| Name | Train | Test | Total | Speedup × |
| --- | --- | --- | --- | --- |
| AIMed | 1500 | 1500 | 3000 | 6.5 |
| BioInfer | 2534 | 2534 | 5068 | 43.6 |
| CC1-LLL | 3785 | 330 | 4115 | 13.1 |
| CC2-IEPA | 3298 | 817 | 4115 | 8.6 |
| CCC3-HPRD50 | 3682 | 433 | 4115 | 4.5 |
| DrugBank | 2472 | 2472 | 4944 | 7.7 |
| MedLine | 635 | 635 | 1270 | 20.7 |

### Baselines

We compare the prediction accuracy and efficiency of FastSK with 3 state-of-the-art string kernel baselines. For DNA and protein data, we baseline against gkmSVM-2.0 [8] and GaKCo [27]. For an NLP string kernel baseline, we use the Blended Spectrum Kernel [12, 11], as it has recently achieved strong results in natural language processing. For FastSK, gkmSVM-2.0, and GaKCo, we perform a grid search over the hyper-parameters *g* ∈ {5, 6, …, 15}, *m* ∈ 0, 1, …, *g* 1, and the SVM margin parameter *C* ∈ {0.001, 0.01, 0.1, 1, 10, 100}. We use 5-fold cross-validation on each training set. For the Blended Spectrum Kernel, we use the authors’ string kernel package ^5^ and use the parameters *k*_*min*_ = 3 and *k*_*max*_ = 5, as recommended by the authors.

We also compare FastSK with bidirectional Long Short-Term Memory (LSTM) memory networks and Character-level CNNs. We select these two straight forward deep learning baselines because we use small-scale datasets. Though models such as AlphaFold or BERT are popular deep learning models for sequence analysis, they require more training data than our datasets contain.

### LSTM Training

To train an LSTM model on each dataset, we perform a grid-search over the embedding and hidden dimensionality ∈ {32, 64, 128, 256}, and number of layers ∈ {1, 2, 3, 4}. We use the Adam optimizer[14] with a learning rate of 0.001. We perform 5-fold cross-validation on each training set. We use the probability scores of the final layer to compute AUC.

### CharCNN Training

To train CharCNN models, we use an architecture with 6 layers; 3 convolutional layers and 3 fully-connected layers.

### SVM Training

To train SVM models using each string kernel method, we use Liblinear [5] via the Scikit-Learn LinearSVC implementation^6^ with L2 regularization. In order to use a kernel method with a linear SVM, we use the empirical kernel map (EKM) strategy. That is, *K* ← *K*_*GSK*_^T^*K*_*GSK*_. To evaluate the SVM models, we use area under the ROC curve (AUC) with Platt Scaling [24] to obtain probabilities. We focus on the AUC metric because many biological sequences datasets are highly imbalanced (very few positive samples in the test set), while a high AUC reflects strong ability to discriminate between true and false positives.

Unless otherwise noted, we run FastSK -Approx with *I*_*max*_ = 50 and *δ* = 0.025. All timing experiments are performed on a server with 12 Intel i7-6850K 3.60GHz CPUs.

### 4.1 Comparing Prediction Performance

#### FastSK achieves excellent prediction performance

Figure 5 shows the AUC results of all baselines across our 27 datasets. Our results prove that FastSK rivals or outperforms previous string kernel algorithms on DNA, protein, and NLP tasks. Moreover, FastSK also outperforms our LSTM and Character-level CNN baselines across all three tasks.

#### 4.1.1 Ablation Study on Approximation Strategy

In this section, we first validate the correctness of our approximation algorithm and then compare our approximation strategy with that of gkmSVM-2.0.

First, our approximation algorithm (FastSK -Approx) is nearly identical to an exact gapped *k*-mer string kernel SVM. As shown in figure 6 (a), a comparison of FastSK -Approx and FastSK -Exact reveals that FastSK -Approx obtains test AUCs only 0.003 points lower than FastSK -Exact. The approximation works so well because, in fact, only a small proportion of the possible mismatch combinations are necessary for excellent performance. We validate this theory in figure 2, as well as figure 12 (appendix).

**Figure 7:**
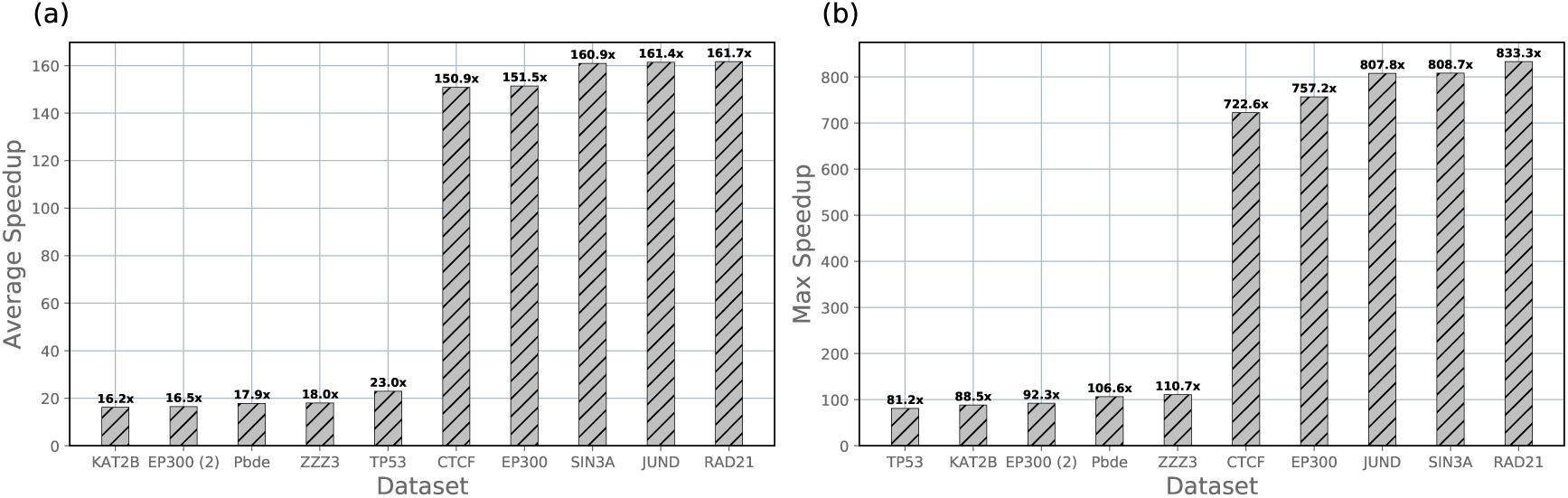
(a) We vary the feature length parameter *g* from 6 to 20 and measure the average speedup achieved by FastSK for each DNA dataset. (b) FastSK achieves maximum speedups of almost 3 orders of magnitude. These are typically achieved when *g* is large.

**Figure 8:**
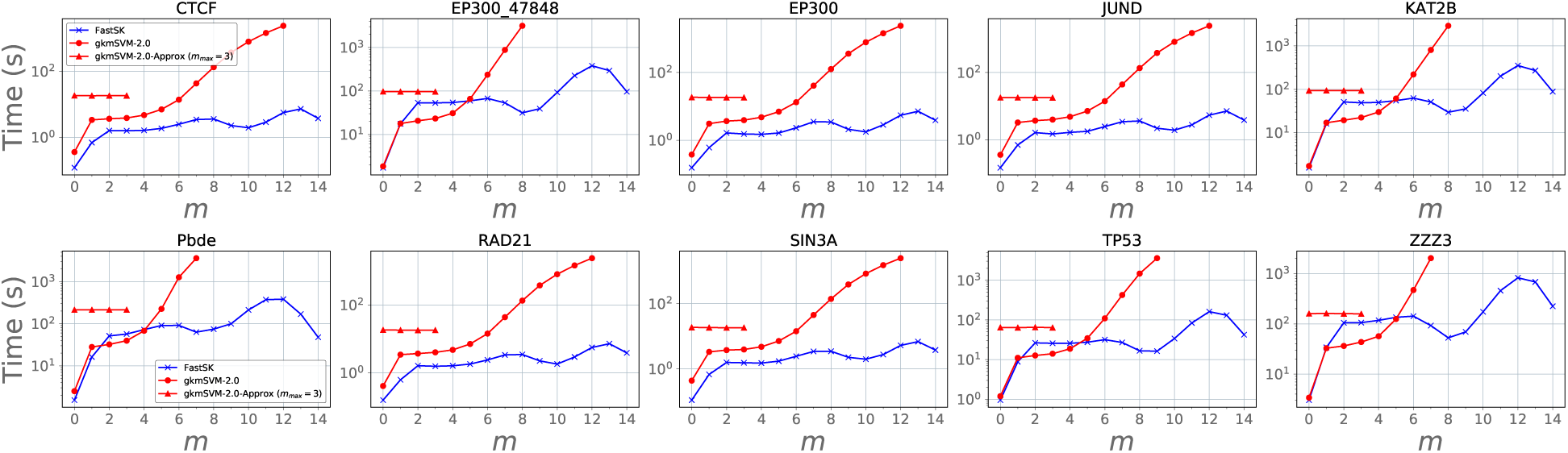
We fix *g* = 16, vary *m* ∈{0, 1, …, 14}, and obtain the kernel computation times for FastSK, gkmSVM-2.0-Exact, and gkmSVM-2.0-Approx. We observe that gkmSVM-2.0-Exact shows exponential slowdowns with respect to *m*, while FastSK remains fast across *m*. Moreover, we point out that gkmSVM-2.0-Approx does not scale to larger to *m >* 3, as it works by limiting the number of mismatch positions to an *m*_*max*_ = 3.

**Figure 9:**
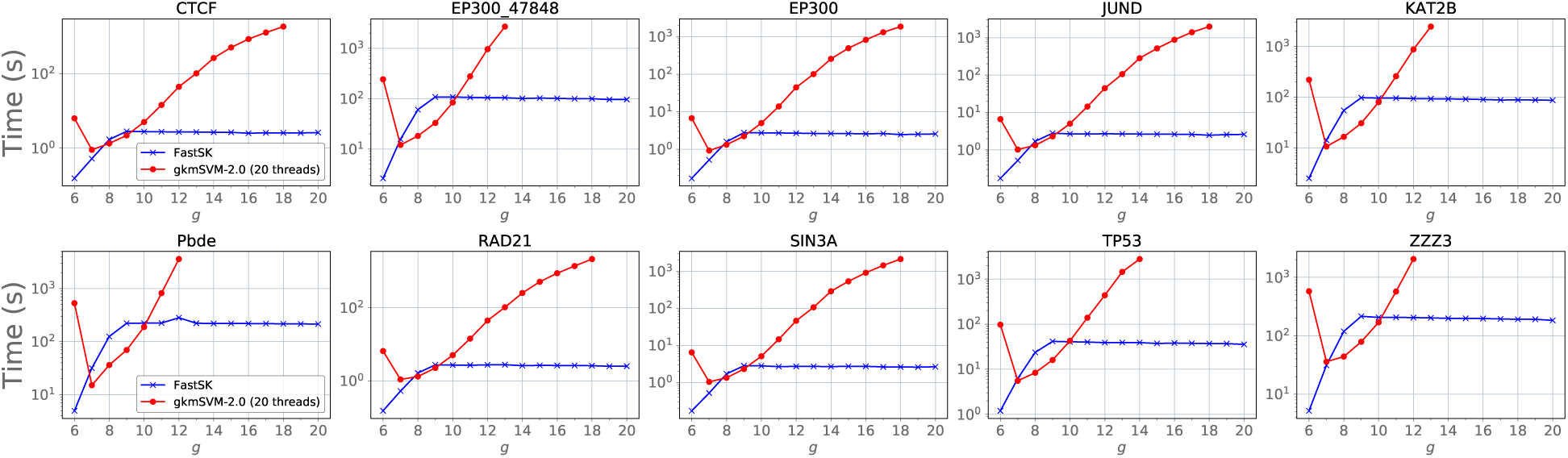
We vary the feature length *g* while keeping the number of informative feature positions fixed at *k* = 6 for each of the 10 DNA TFBS datasets. We run FastSK with the maximum number of mismatch combinations set to *I*_*max*_ = 50 and run gkmSVM-2.0 using 20 threads. We stop each algorithm early once the kernel computation time exceeds 1800s (30 minutes). We demonstrate that FastSK is effectively *O*(1) with respect to the feature length *g*.

**Figure 10:**
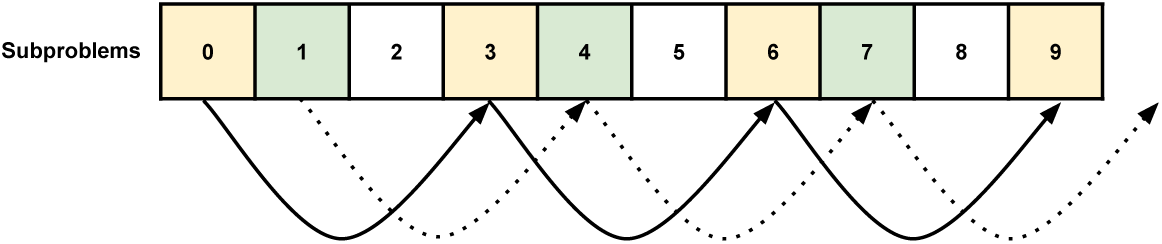
An illustration of how subproblems (partial kernel matrices *P*_*i*_) are divided between threads. If there are 9 mismatch position combinations and *t* = 3 threads, then thread 0 (solid arrow and yellow cells) handles the combinations/subproblems 0, 3, 6, and 9. After finishing each subproblem, it aggregates the results into a local sub-sum matrix and skips forward *t* positions. The dashed arrow (green cells) shows the procedure for thread 1. Once each thread has run out of subproblems to handle, it proceeds to aggregate its results into the full kernel matrix *K*.

**Figure 11:**
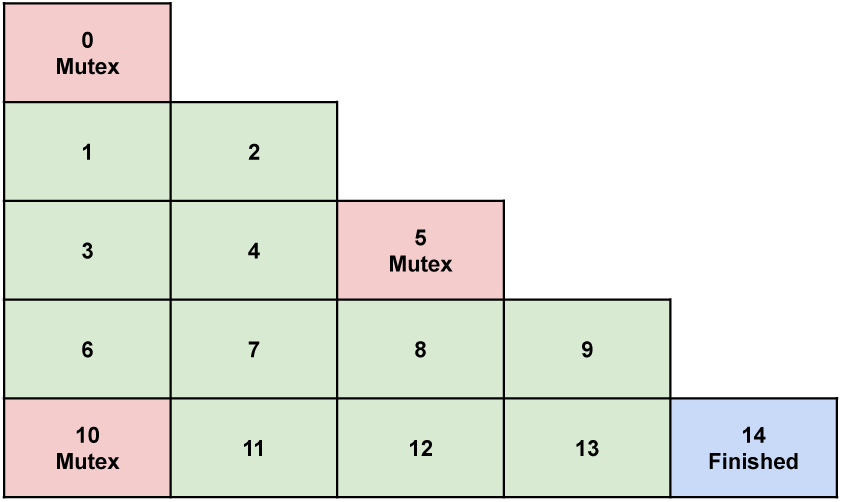
An example of our matrix aggregation and synchronization scheme using the lower triangle of a kernel matrix. The indexes show the order in which cells are accessed by the threads. Each time a thread reaches one of the red cells (labeled “Mutex”), it locks the indices up until the next mutex location. When it reaches the next mutex location, it unlocks the previous memory region to permit a new thread to enter. It also locks the next memory region. Though shown here as a triangular matrix, the kernel matrix is actually a single contiguous array in practice. This maximizes cache performance.

**Figure 12:**
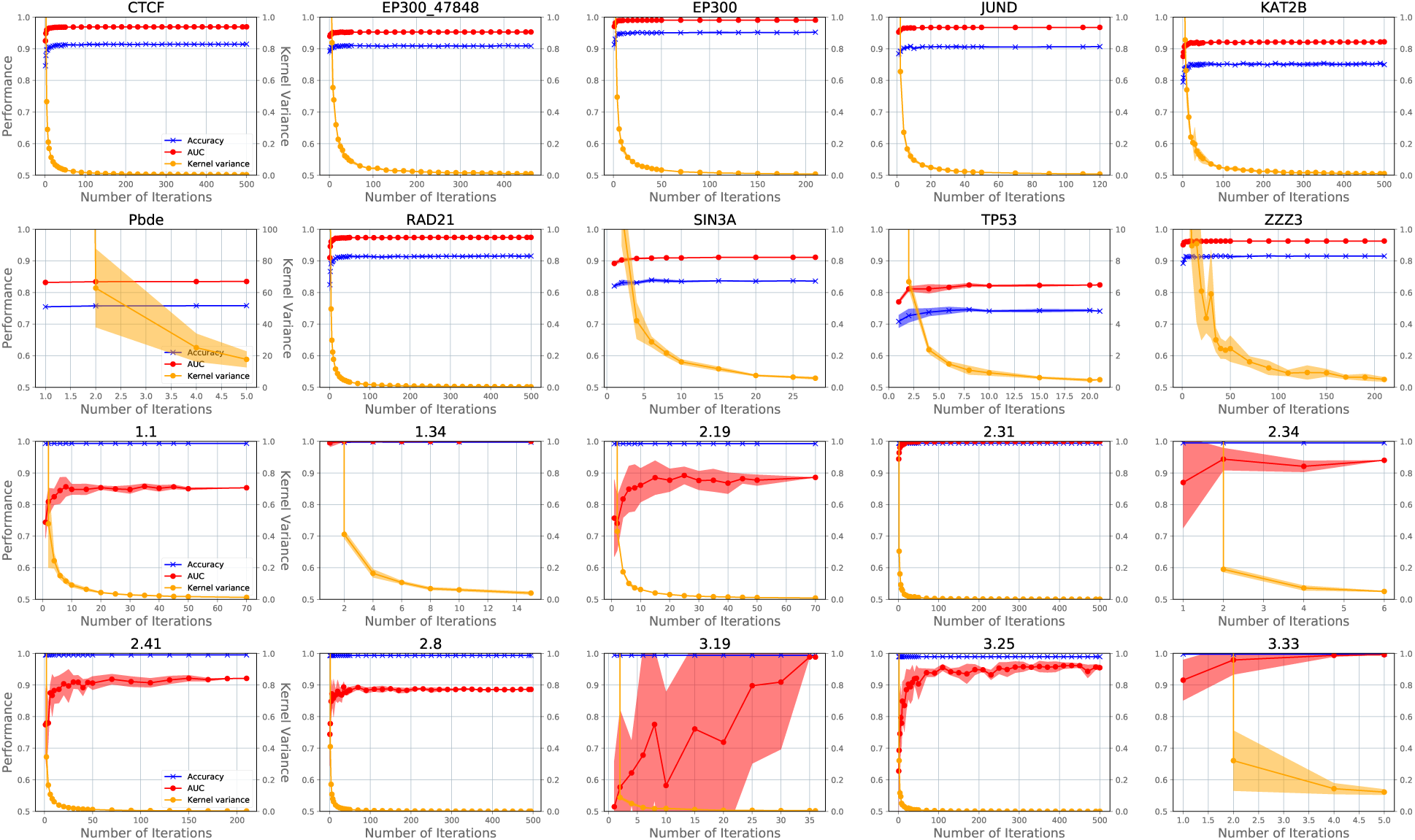
We vary the maximum number of mismatch positions sampled (determined by the *I*_*max*_ parameter) used for kernel computation. We then train and test an SVM and show the resultant test accuracy and AUC. These results show that extremely few mismatch positions for kernel computation usually results in poor test performance, but that test performance increases with the mismatch positions sampled up until several dozen. This finding provides insight into why FastSK -Approx works so well. However, these results also showcase an unexpected property of gapped *k*-mer SVMs: in some cases extremely few mismatch combinations are actually necessary for excellent performance. We propose exploring this phenomenon as a direction for future research.

Second, we show that our approximation algorithm is favorable compared to that of gkmSVM. Figure 6 (b) shows that FastSK-Approx consistently maintains excellent AUCs and low kernel timings. Meanwhile, we show that gkmSVM-Approx is actually optimized for kernel computations that are *already* fast; as the *g* parameter increases, gkmSVM-Approx both slows down and dramatically drops off in AUC. It therefore works best when *g* is small, but these are precisely the cases where an approximation algorithm is unnecessary. Figure 6 (c) further illustrates this problem: as *g* increases, gkmSVM-Approx’s AUCs deteriorate rapidly. In contrast, FastSK’s AUCs remain quite stable. We therefore conclude that FastSK-Approx is both fast across the parameter space and produces reliable results. These conclusions are further explored in section 4.2.

### 4.2 Comparing Speed and Scalability

In this section, we experimentally analyze the scalability of FastSK and compare with the primary baseline, gkmSVM-2.0. First, we show that on average FastSK is between 1 and 3 orders of magnitude faster than gkmSVM-2.0. Second, we show that FastSK easily scales to larger numbers of mismatches *m*, while gkmSVM-2.0 does not. Third, we show that FastSK easily scales to larger feature lengths *g*, while gkmSVM-2.0 does not. Taken together, these results demonstrate a major contribution of this work: unlike previous works, FastSK is practical and easily navigates the gapped *k*-mer parameter space of *g* and *m* values.

#### FastSK is orders of magnitude faster

As illustrated in table 1, tables 2 and 3 (appendix), and figure 7, FastSK achieves large speedups over the baselines. We vary the feature length *g* and time kernel computation for each value. This simulates the necessary process of searching the full space of parameters in a grid search. We show that on average, FastSK is hundreds of times faster than gkmSVM. Maximum speedups are over 800×.

We evaluate FastSK using 10 DNA sequence based transcription factor binding site classification datasets (downloaded from the DeepBind [1]). We measure the factor of speedup in kernel computation time relative to gkmSVM-2.0, showing that FastSK is typically hundreds of times faster than gkmSVM-2.0. Specifically, for each dataset, we vary *g* from 6 to 20, while fixing *k* = 6, and then time the kernel computations to obtain the speedups. We run FastSK -Approx with *I*_*max*_ = 50, as was used to obtain the results shown in figures 5 and 6. Timing figures for all 10 DNA datasets are shown in figure 9

#### 4.2.1 Scalability Analysis When Varying Mismatches *m* and Feature length *g*

**FastSK scales in** *m*We show that FastSK scales more efficiently than gkmSVM-2.0 with respect to the mismatch parameter *m*. Figure 8 shows gkmSVM-2.0 is exponential in *m*, as discussed in section 3.5. On the other hand, gkmSVM-Approx attempts to circumvent this problem by using a parameter *m*_*max*_ = 3. However, this has the downside of severely curtailing the size of the parameter space. The decision comes at the expense of flexibility and performance, as we showed in figure 6. The approximation algorithm used by FastSK overcomes these challenges.

**FastSK scales in** *g*We show in figure 9 that gkmSVM-2.0 grows exponentially with the feature length *g*, while FastSK is effectively 𝒪(1) with respect to *g*. Moreover, FastSK does not exhibit dramatic AUC decay as *g* increases. Therefore, FastSK is able to span the gapped *k*-mer parameter space.

## 5 Conclusion

In this work we introduced a fast and scalable string kernel SVM algorithm called FastSK. FastSK rivals state-of-the-art string kernel SVMs in test performance, while running 1-3 orders of magnitude faster on average. Unlike previous methods, FastSK is scalable with respect to the *g* and *m* gapped *k*-mer parameters. Moreover, it outperforms two deep learning baselines. FastSK makes three high-level contributions:

1. Across 10 DNA based transcription factor binding site (TFBS) prediction datasets, FastSK consistently matches or outperforms the state-of-the-art gkm-SVM-2.0 algorithms in AUC, while achieving average speedups in kernel computation of ∼ 100*×* and speedups of ∼ 800*×* for large feature lengths.
2. We showed that FastSK outperforms character-level recurrent and convolutional neural networks across all 10 DNA tasks.
3. We further demonstrated the utility and robustness of FastSK by testing it on 10 protein remote homology detection tasks and 7 English-languaged medical named entity recognition tasks. We showed FastSK outperformed the baselines across all 17 datasets.

## 6 Appendix

### 6.1 More Details for Implementation

#### Multithreading

As shown in equation 12, we decompose the kernel function into a summation of 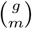 partial kernel matrices *P*_*i*_, with each *P*_*i*_ holding the gkmer counts for a single combination of mismatch positions. Observing that each *P*_*i*_ is a completely independent subproblem, we exploit the decomposition to create a multithreaded implementation.

To map threads to subproblems, we create a “work queue”—a length 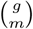 array, where each element indicates a mismatch combination subproblem to solve. If there are *t* threads (a user-specified argument), we divide the work queue into *t* equally sized portions. In practice, each thread simply moves forward *t* positions in the work queue when it completes a subproblem. An illustration is shown in figure 10. A thread’s local results are aggregated into its own temporary matrix; since all *P*_*i*_ are added together in the end, each thread simply aggregates its partial results as it finishes them. Then once each thread finishes, it adds its partial results into the full kernel matrix *K*.

#### Synchronization

Once each thread finishes computing its subset of the 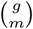 partial kernel matrices, these are aggregated into the kernel matrix *K* as per equation 12. But because each thread must access the same *K* in memory, we must obviate potential race conditions. To ensure synchronization and maximize the number of threads able to access *K* simultaneously, we use a set of mutex locks. Intuitively, the idea is to create a set of evenly sized memory regions such that only one thread is allowed in each region at a time; a thread enters a region, locks it, and performs its aggregations. When it finishes it unlocks the previous regions, permitting a new thread to enter, and then locks the next region. An illustration using a small kernel matrix is shown in figure 11.

### 6.2 Protein and NLP Dataset Details and Speedups

We evaluate FastSK using 10 SCOP project protein remote homology detection datasets and 7 medical named entity recognition datasets. We show the kernel computation time factor of speedup achieved by FastSK over GaKCo and Blended Spectrum on each dataset. The average factors of speedup are 94.7× and 15.0×, respectively.

### 6.3 More Experimental Results

#### 6.3.1 More Ablation Analysis when Varying approximation parameter I

**Figure 13:**
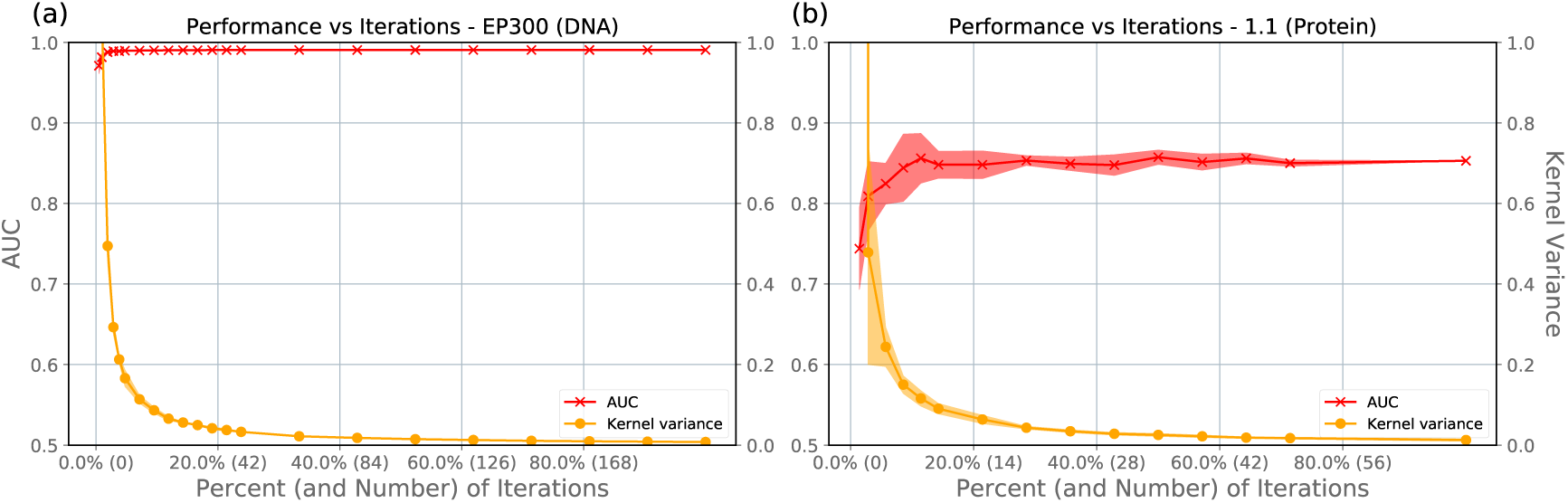
We vary the number of iterations (i.e., the number of mismatch combinations sampled) used by the approximation algorithm to compute the kernel matrix. The x-axis shows the percent of the maximum number of iterations (determined by the optimal *g* and *m* parameters for each dataset). In addition, it shows the number of iterations used. The left y-axes show the AUC obtained after each number of iterations. The right y-axes shows the average variance of the kernel matrix. These results confirm our hypothesis that the approximation algorithm converges rapidly with the number of iterations and shows that very few iterations are needed for excellent test AUC. Each point shows the average of 5 runs of the approximation algorithm, with the shading indicating 95% confidence interval. These results are shown for: (a) The EP300 TFBS dataset (DNA classification) and (b) the 1.1 protein classification dataset.

#### 6.3.2 More Scalability Analysis when Varying m

**Figure 14:**
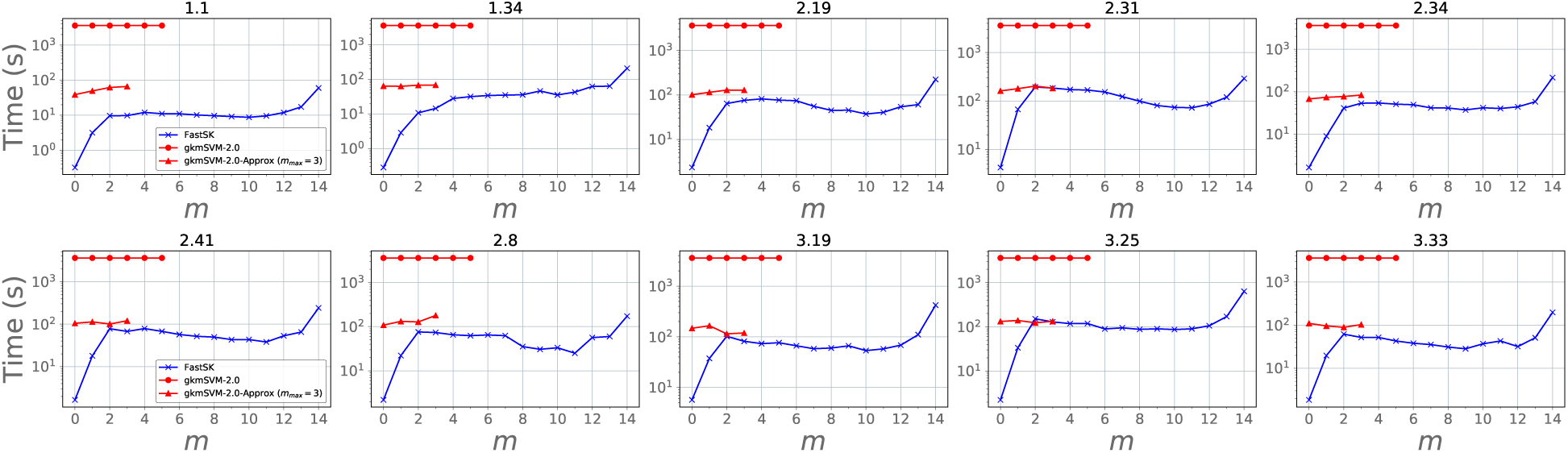
Across each of the protein datasets, we fix *g* = 16 and vary *m* ∈ {0, 1, …, 14}. All gkmSVM-2.0-Exact experiments took over an hour and timed out. As noted in figure 8, the gkmSVM approximation algorithm does not work for *m >* 3. Therefore, we argue FastSK efficiently scales across the parameter space for both DNA and protein classification tasks.

#### 6.3.3 More Scalability Analysis when Varying g

**Figure 15:**
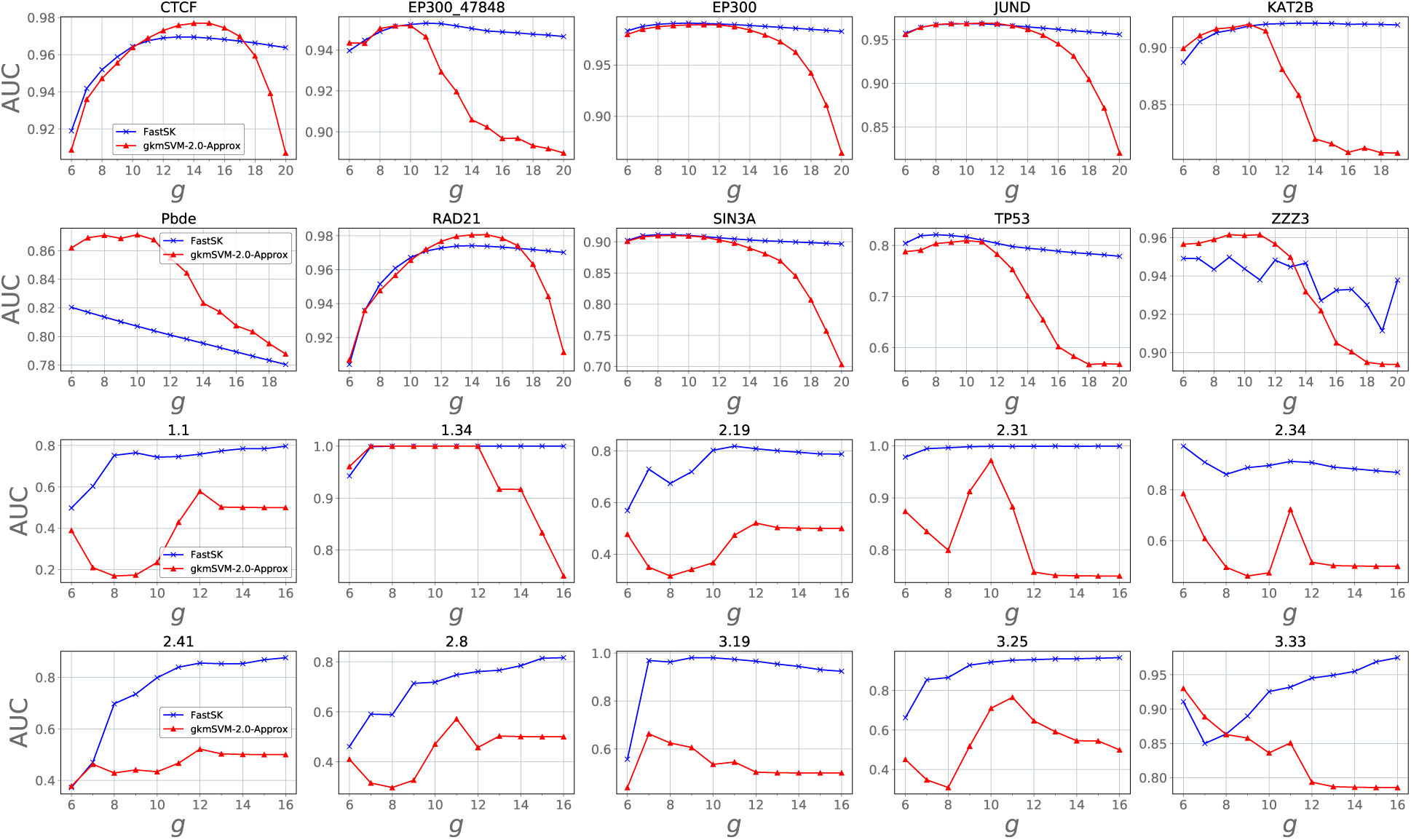
We explore how test AUC changes with varying *g* for FastSK and gkmSVM-2.0 across DNA and protein datasets. Here we show that optimal AUC is often obtained when *g* is large. Moreover, we note that even in cases when large *g* is suboptimal, those values must still be searched in a thorough grid search. Though FastSK easily scales to large values of *g*, gkmSVM-2.0 is unable to do so, as its AUC values rapidly degrade as *g* increases. Note: we use gkmSVM-Approx in this figure, as using their exact algorithm is impractical, as figure 9 makes clear.

#### 6.3.4 Multithreading

In this section we examine how kernel computation time is a function of the number of threads for FastSK -Exact, FastSK -Approx, and gkmSVM-2.0. To start, we point out that FastSK -Approx generally does not benefit from using multiple threads. There are two reasons. First, our primary implementation of the multithreaded FastSK -Approx algorithm improves approximation quality, not speed. This is because each thread simply iterates independently until it converges. After all threads converge, their mean kernels are aggregated and normalized. Nothing about this improves speed per se. Second, we did create an option to *accelerate* FastSK -Approx, but the speedups were marginal. This option is identical to FastSK -Exact, except with the *I*_*max*_ parameter set to a user-specified value. That is, it simply skips the variance and mean kernel matrix computations, using *t* threads to compute *I*_*max*_ partial kernels *P*_*i*_ in total. In other words, each thread computes ≈*I*_*max*_*/t* partial kernels. However, we have shown *I*_*max*_ ≈50 is usually sufficient. This is such a small number of iterations that it does not stand to benefit much from multithreading. In fact, the synchronization overhead in use-cases like this typically negate the benefits of multithreading.

That being said, FastSK -Approx with just *one* thread is still vastly faster than either FastSK -Exact or gkmSVM-2.0 with *twenty* threads. Unlike FastSK -Approx, these algorithms benefit greatly from more threads, as we show. But nevertheless, FastSK -Approx is much faster. We claim this is one of the key merits of this work: fast kernel computation without multithreading at all. Yet even in the case of exact kernel computation, we claim another key merit: at each number of threads FastSK -Exact is significantly faster than gkmSVM-2.0, which we show in 16.

##### FastSK outperforms gkmSVM-2.0 for any number of threads

As shown in figure 16, FastSK -Exact is substantially faster than gkm-SVM2.0 for each number of threads. FastSK -Approx is vastly faster still, even when using just 1 thread. We show similar behavior across all 10 DNA datasets in the appendix in figure 16.

**Figure 16:**
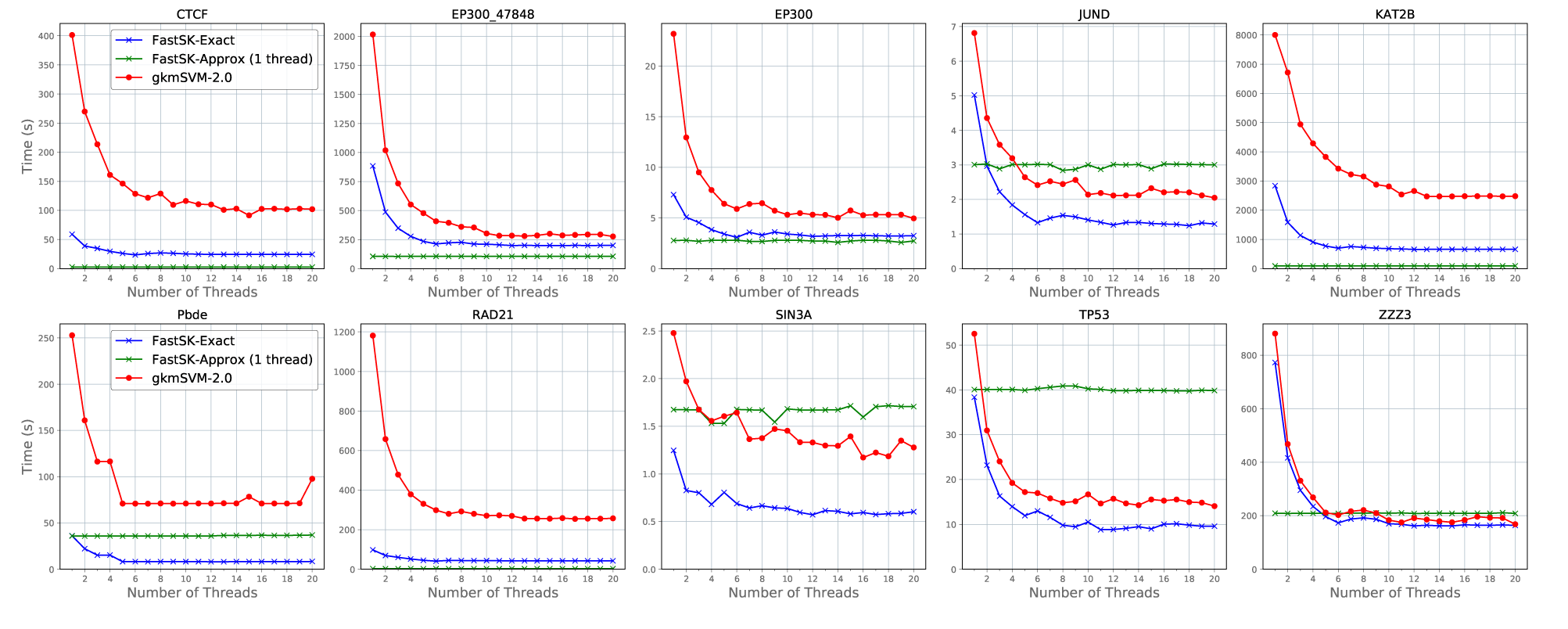
We vary the number of threads used for kernel computation across all 10 DNA datasets. Each kernel is computed using the optimal parameters.

##### FastSK is better for multithreading than GaKCo or Blended Spectrum

Neither GaKCo nor the Blended Spectrum kernel make adequate use of multithreading. GaKCo limits the number of possible threads to *m*, while Blended Spectrum only allows one thread.

### 6.4 Future Directions

We identify several promising areas where future work could be directed.

1. Finding even simpler gapped *k*-mer formulations. The success of FastSK -Approx and the curiously small number of mismatch combinations needed in many cases suggest simpler exact algorithms could exist. The goal here would be to shed the 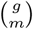 coefficient from the exact algorithm. An intuition is that it seems only a few mismatch positions are necessary to obtain most or all of the *unique* gapped *k*-mers and that the number of unique gapped *k*-mers is what ultimately matters.
2. Reducing memory usage. We currently use 𝒪 (*n*^2^) memory to store the kernel matrix. This is intractable for *n* greater than several tens of thousands. Future work should implement techniques to avoid storing the full kernel matrix in memory at once. For example, [18] only computes small batches of the kernel matrix at a time, greatly reducing the memory footprint.
3. Capitalizing on interpretability. String kernel methods have interpretable features and many works have identified the important features from string kernel methods [26, 19]. Future work can use methods such as Data Shapley [9] to analyze the most salient features. The fact that so few features appear to be necessary in our results suggests a small number of features are critical, while most are disposable. Future interpretability work could study this hypothesis.
4. Low rank approximations of the gram matrix *K*. The efficiency of kernelized SVMs can be improved via the Nyström method [31], which creates a low-rank approximation of the kernel matrix. This method still requires 𝒪(*n*^2^) space for the kernel matrix *K*, but it ultimately provides an approximation 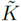 to be used by the SVM optimizer. For inspiration, [32] shows a linear SVM method inspired by Nyström’s method.

### Footnotes

1 Available for download at https://github.com/Qdata/FastSK/. Install with the command make or pip install

2 We hereafter use “gap” and “mismatch” interchangeably.

3 Available at https://github.com/QData/FastSK or with the command “pip install fastsk”

4 https://github.com/pybind/pybind11

5 Available for download at: http://string-kernels.herokuapp.com/

6 https://scikit-learn.org/stable/modules/generated/sklearn.svm.LinearSVC.html

